# Protocol for the Synthesis of Green Fluorescence Carbon Quantum Dots and Biological Application

**DOI:** 10.1101/2025.10.28.685185

**Authors:** Himanshu Khatri, Ankesh Kumar, Raghu Solanki, Pankaj Yadav, Dhiraj Bhatia, Amit K. Yadav

## Abstract

Carbon quantum dots (CQDs) are materials that are made up of particles that are only a few nanometers wide. They have no dimensions and can change their fluorescence qualities. They are also verybiocompatible and have a lot of potential to change the chemistry of their surface. Their eco-friendly synthesis from precursors, like citric acid (CA) and ascorbic acid (AA), provides a sustainable approach to highly fluorescent nanomaterials with promising applications in bioimaging, drug delivery, and other biomedical fields. Green-emitting carbon quantum dots (GCQDs) were synthesized via a simple reflux approach using citric acid and ascorbic acid as biomolecular precursors in a solvent solution of ethanol-water (1:2). The obtained GCQDs solution was purified using a dialysis process. Besides this, pH maintenance, lyophilization, and further submission for detailed physicochemical evaluation and characterization are done. The spectroscopic analysis, including UV–visible spectroscopy, fluorescence spectroscopy, and FTIR, confirmed the presence of surface functional groups and strong fluorescence properties and quantum yield calculations of the synthesized GCQDs. Microscopic and structural analyses using XRD, AFM, and TEM revealed the nanoscale, predominantly spherical morphology of the synthesized GCQDs. In addition, in vitro biological evaluations such as MTT assay and cellular uptake analysis were undertaken to evaluate cytocompatibility and intracellular distribution. We study the endocytosis pathway of these GCQDs, with size variations ranging from 3 to 5 nm, in mouse tissue-derived primary cells, tissues, and zebrafish embryos. GCQDs were internalized into mouse kidney and liver primary cells through a clathrin-mediated pathway. The findings confirmed that the produced GCQDs exhibit good water solubility, favorable biocompatibility, and significant potential as candidates for biomedical imaging applications.

## 1. Introduction

Carbon quantum dots (CQDs) are unique multifunctional nanomaterials characterized by properties such as strong fluorescence, ultra-small nanoscale dimensions, versatile surface chemistry, excellent photostability, and minimal toxicity. These features make them highly suitable for a wide range of biomedical applications, including early tumor detection, bioimaging, photoacoustic imaging, drug delivery, and biosensing. In addition to their biomedical potential, CQDs are environmentally friendly, cost-efficient, and exhibit high electrical conductivity. Structurally, they are zero-dimensional materials, typically below 10 nanometres in size. Owing to these advantages, CQDs are considered promising candidates in the treatment of several diseases, such as cancer, infectious illnesses, ophthalmic disorders, neurological conditions, and cardiovascular diseases^1–3^.

Green synthesis of CQDs is gaining increasing attention in nanotechnology, with natural products being widely explored as precursors. Researchers favor these resources because they are cost-effective, eco-friendly, easy to handle, and abundantly available. Among them, plant-based materials such as fruits, peels, leaves, vegetables, and other botanical derivatives are the most commonly used sources for producing fluorescent CQDs^4^ ^5,6^.

Citric acid is one of the most common weak organic acids, with a well-liked carbon precursor due to its good biocompatibility, low toxicity, and cost-effectiveness. These citric acid-based CQDs not only have a high Quantum yield but also show great photoluminescence from blue regions to red regions^7,8^.The most common sources are orange juice & lemon juice, which are popular for the synthesis of fluorescent CQDs^9,10^. Ascorbic acid is a reduced form of vitamin C, which is synthesized from the hexose sugar(D-sorbitol plays a fermentation step, and Bioconversion takes place into L-ascorbic acid) & it occurs in only three major isomeric forms, which are L-ascorbic acid, L-arabo-ascorbic acid, and D-arabo-ascorbic acid. From this, L-ascorbic acid is widely used. Ascorbic acid is synthesized by all higher category plant species^11,12^.

Green-emitting carbon quantum dots (GCQDs), were synthesized from biocompatible precursors such as citric acid (CA) and ascorbic acid (AA), providing a sustainable approach for biomedical applications. Following synthesis, the GCQDs were carefully characterized by advanced instrumentation to understand their structural, physicochemical, and optical features, which also helps in quantum yield calculations. Transmission electron microscopy (TEM) and atomic force microscopy (AFM) were used to determine particle size and morphology of the GCQDs, while X-ray diffraction (XRD) confirmed the crystalline nature of these GCQDs. Optical behavior was assessed through UV–Vis absorption and fluorescence spectroscopy, and quantum yield (QY) measurements. Quantum yields were calculated by taking standards as Rhodamine B (RB) and fluorescein isothiocyanate (FITC). Systematic study of the biological performance of GCQDs was then evaluated in a stepwise manner. MTT cytotoxicity assays in cultured cells demonstrated low toxicity. Cellular uptake and inhibition studies in primary kidney and liver cells showed that GCQDs enter cells via both clathrin-mediated and clathrin-independent pathways. Ex vivo experiments with mouse kidney tissue revealed a concentration-dependent penetration of GCQDs, while in vivo studies using zebrafish larvae (72 hpf) according to OECD Guidelines confirmed safe, dose and time-dependent biodistribution without any developmental abnormalities. Overall, these findings bring together synthesis, characterization, and biological assessment, underscoring the potential of GCQDs as safe, versatile nanoprobes for bioimaging and future applications in nanomedicine.

## 2. Applications of Carbon Quantum Dots

### Chemical sensing

There has been a significant surge in interest in recent decades to study the potential of carbon nanodots, since they are inexpensive, compatible, and sensitive nanosensors for chemical sensing. This is often discovered by continuous monitoring of changes in fluorescence characteristics or absorbance in the presence of a certain analyte. the existence of heteroatoms might show greater potential towards the sensing performance, and they are customized to interact with the particular analyte^13–15^. These carbon quantum dot band sensors are employed in human serum, which is used to identify and boost their usefulness for periodic checks and monitoring of zoledronic acids^16^.

### Bioimaging

Bioimaging is one of the most explored uses of CDs owing to their typically low cytotoxicity and resilience to photobleaching^13,17^. Carbon dots (CDs) have emerged as attractive nanomaterials for bioimaging applications owing to their low cytotoxicity and great biocompatibility^18,19^. They may rapidly infiltrate cells and are largely dispersed inside the cytoplasm. CDs have been effectively exploited for imaging in numerous biological systems, including cancer cells, microalgae, zebrafish, murine organs, and even for non-biological applications like as fingerprint identification. However, their metabolic pathways and elimination routes in vivo remain inadequately characterized, underscoring the need for greater exploration into their biodistribution and clearance processes^20,21^.

### Biomedical Applications

Carbon quantum dots (CQDs) possess unique physicochemical and catalytic properties that make them highly suitable for biomedical applications^22–24^. Their small size, coupled with excellent biocompatibility, offers significant potential for use in drug delivery systems. In addition, CQDs exhibit very low toxicity, high hydrophilicity, and remarkable water solubility. Their wavelength-dependent photoluminescence emission, along with strong chemical stability, further enhances their versatility, making them valuable candidates for a wide range of biomedical applications^25–29^.

## 3. Overview of the procedure

Green carbon quantum dots were manufactured using citric acid and ascorbic acid as natural precursors via a reflux-assisted approach. The method began with thorough sterilization and cleaning of glassware, followed by dissolving the precursors in an ethanol–water mixture (1:2), vortexing, and sonication to achieve homogeneity. The mixture was then refluxed at 130 °C for 12 hours under constant stirring. After completion, the reaction product was cooled, neutralized to pH 7, and filtered by 24-hour dialysis with occasional water replacement. Finally, the purified GCQD solution was frozen at −80 °C and lyophilized to obtain dry powdered nanomaterial for future characterization and biological evaluation.

### 3.1 Limitations of the protocol

- **Energy and Time-Intensive Process:** The reflux-assisted synthesis protocol involves prolonged heating (≈approximately 12 hours), followed by extended purification via dialysis (24 hours) and lyophilization (24 hours). Such multi-step procedures are not only energy-intensive but also time-consuming, reducing overall throughput and efficiency.
- **Low Production Yield:** Despite successful synthesis, the overall production yield of GCQDs remains relatively low, which limits the availability of sufficient quantities for large-scale biomedical or industrial applications.

### 3.2 Applications of the protocol

The reflux method is widely used for the synthesis of CQDs because of their simple, cost-effective, aqueous-based nature and their suitability for producing highly stable fluorescent CQDs^30,31^. This reflux method involves the slow and low-temperature heating under complete condensation, which minimizes the solvent evaporation and ensures that no solution loss takes place. This reaction method allows for stable reaction conditions for a very long duration of time, making them suitable for the synthesis of carbon quantum dots^32^. In this reflux method, we further perform dialysis, which is meant to remove the unbound chemical entities, and we get a uniform size distribution, which is directly helpful for cellular uptake studies and bioimaging^33,34^. These reflux method of synthesis gives a good quantum yield and dry crystalline powder of GCQDs^35^.

### 3.3 Comparison with other methods

#### Hydrothermal method

Hydrothermal pressure requires high temperature and pressure, through which we get uniform and crystalline CQDs, but needs an autoclave and is less scalable, while the reflux method requires moderate temperature and normal pressure. The reflux method is simpler and safer than the hydrothermal method and produces amorphous CQDs with broader size variation. Both methods yield functional fluorescent CQDs, but the choice depends on whether you prioritize ease & scalability (Reflux) or high crystallinity & stability(hydrothermal)^36,37^.

#### Microwave-Assisted method

Microwave-assisted synthesis provides rapid and volumetric heating, produces bright and smaller CQDs within a few minutes, and has a high quantum yield. however, this method often requires specialized equipment with its operators, and they may also yield batch-to-batch variations. While reflux synthesis employs gentle, prolonged heating at atmospheric pressure over extended periods, it offers simple equipment, reproducible conditions, and safe handling^38,39^.

## 4. Materials Required

### 4.1 General Chemicals

1. Citric acid Anhydrous (SRL chemicals),
2. Ascorbic acid 99.5% (Loba chemicals),
3. Deionized water (From Milli-Q direct water purification system, Merck Millipore),
4. Ethanol 99.5%v/v (Changshu Hongsheng Fine Chemicals Co., Ltd).
5. Rhodamine B
6. FITC (Fluorescein Isothiocyanate) (Sigma Aldrich)

### 4.2 Cell culture

1. Dimethyl sulfoxide (DMSO; Himedia)
2. Phosphate buffer saline (PBS; Gibco)
3. Trypsin-EDTA (Thermo Fisher Scientific)
4. Dulbecco’s Modified Eagle Medium (DMEM; Gibco)
5. Fetal bovine serum (FBS; Sigma-Aldrich)
6. DAPI
7. MTT salt
8. 4% paraformaldehyde
9. 1x collagenase D
10. DMEM: F12
11. Transferrin
12. Galactine-3
13. Pitstop-2
14. Lactose
15. E3 media
16. Liquid nitrogen

### 4.3 Equipment

1. Pipettes (Eppendorf)
2. Water bath (scientiz,cat.no.SB800DT)
3. Pipette tips (Tarsons)
4. Refrigerated benchtop centrifuge(Eppendorf,cat.no.5810R)
5. Ice machine (Coolium, cat. no. FM100)
6. Ultra-pure water purifier (Milli-Q, cat. no. C85358)
7. Freezers and refrigerators (Haier, cat. no. BCD-149WDPV; Haier, cat. no. DW-251262; Haier, cat. no. DW-861.626)
8. pH meter (Mettler Toledo, cat. no. 210-S)
9. Bath. Sonicator (Cole-Parmer, cat. no. 08849-00
10. Vortex mixer (Thermo Scientific, cat. no. 88880018)
11. Autoclave
12. Dessicator
13. Magnetic stirrer
14. Analytical balance
15. Microscope (Nikon)
16. Confocal scanning microscope ((Leica Sp8))
17. Fluorescence Spectrophotometer
18. UV-Visible spectrophotometer
19. UV-VIS/Fluorescence cuvettes
20. Mica substrate
21. Wash bottle (100 mL)
22. Centrifuge tubes (Falcon 15 mL & 50 mL)
23. Parafilm tape
24. Glass vials
25. Snakeskin dialysis membrane 3.5K (Thermoscientific cat.no.AC403628)
26. Water pump system
27. Magnetic bead
28. Syringe needle
29. Hot air oven

### 4.4 Cell Culture

1. Hemacytometer (Thermo Fisher Scientific)
2. 12 mm Coverslips (Bluestar)
3. Microscopic Slides (Bluestar)
4. Cell culture microscope (Carl Zeiss)
5. Micro centrifuge tubes (100 µL,200 µL,1 mL,2 mL)
6. High-resolution Transmission electron microscope
7. Cell culture Incubator (Thermo Fisher Scientific)
8. 4-well plate (Sigma Aldrich)
9. 96-well plate(Sigma Aldrich)
10. Serological pipettes (Tarsons)
11. Cell culture flask, 25 cm^2^ (NEST)
12. Cell culture dish, 10cm (NEST)
13. Multichannel Pipettes (Thermo Fisher Scientific, cat. no.25300-62)
14. Scalpel

### 4.5 Glassware

1. Silicon oil bath
2. RBF holder
3. Beaker (100 mL,250 mL,500 mL,1000 mL)
4. Round-bottom flask 100 mL
5. Measuring cylinder
6. Reflux condenser

**Note: -** All chemicals should be handled after wearing personal protective equipment, such as a lab coat and gloves.

**Note: -** All Chemicals should be used according to the manufacturer’s protocol.

## 5 Reagent preparation

### 5.1 10 M NaOH preparation

- Weigh accurately 4 g NaOH pellets and add 10mL of water, and cool down for a few minutes.

**Note: -** Use a heat-resistant beaker because it will heat up due to an exothermic reaction.

### 5.2 Aqua Regia

- Firstly, turn on the fume hood and place a 250 mL beaker
- Add 135mL of HCL and slowly add 45mL of Nitric acid to it (Nitric acid: HCL 1:3) **Note: -** All chemicals should be handled after wearing personal protective equipment, such as a lab coat, face mask, and gloves. Aqua Regia is extremely corrosive and can cause severe burns. It also releases toxic chlorine (Cl) and nitrogen oxides (NOx) fumes.

**Note: -** Always prepare immediately before use. Do not store aqua regia, as it decomposes and may cause container rupture. Preparation and use must be carried out in a certified chemical fume hood.

### 5.3 1X Phosphate-buffered saline (PBS)

- Dissolve 8.0 g NaCl, 0.2 g KCl, 1.44 g Na2HPO4, and 0.24 g KH2PO4 in 800 mL deionized water, and adjust the final pH to 7.4 using HCl.
- Add deionized water to bring the solution volume up to 1,000 mL. Filter-sterilize (0.22 µm), and store PBS solution at 4°C for no more than 6 months.

### 5.4 MTT (3-(4,5-dimethylthiazol-2-yl)-2,5-diphenyltetrazolium bromide) tetrazolium solution

- Gently dissolve MTT in Dulbecco’s Phosphate Buffered Saline, pH=7.4 (DPBS) to 5 mg/ml.
- Filter-sterilize the MTT solution through a 0.2 µM filter into a sterile, light-protected container.
- Store the MTT solution, protected from direct light, at 4°C for frequent use or at - 20°C for long-term storage.

### 5.5 FITC(fluorescein isothiocyanate)

- Directly dissolve the 50µL of FITC in 1 mL of DMSO and mix using a vortex mixer.

### 5.6 Rhodamine B (10 µ/mL)

- Dissolve 1mg rhodamine in 1 mL milli-Q. further take 0.1 mL from the 1 mg/mL solution and dissolve in 9.9mL of milli-Q water. The final concentration is 10 µL.

### 5.7 1× Collagenase D: -

- Weigh out 10 mg Collagenase D. Dissolve in 10 mL sterile PBS (without Ca²/Mg²). Filter-sterilize and keep on −20 °C until use.

### 5.8 DMEM Solution

- Add 450 mL DMEM (glutamine base) to an autoclaved bottle and add the supplement with 50 mL FBS (stored at –80 °C).
- Add 1× Penicillin-Streptomycin, HEPES, sodium bicarbonate, and sodium pyruvate to prepare the solvent control medium.
- Filter-sterilize the complete medium into the autoclaved bottle and store at 4 °C.

### 5.9 Transferrin solution

- Accurately weigh 5mg of transferrin and directly dissolve it in 1 mL of Milli-Q water, using a vortex mixer to ensure even dissolution.

### 5.10 Galactine solution

- Accurately weigh 5mg of galactine and directly dissolve it in 1 mL of Milli-Q water, using a vortex mixer to ensure even dissolution.

### 5.11 Lactose stock solution

- To prepare 1 mL of 1 M, dissolve 342.3 mg lactose(molecular weight of lactose monohydrate 342.3 g/mol) in 1.0 mL sterile water.

### 5.12 Pitstop-2 stock solution

- To prepare 1 mL of the stock solution, we have to dissolve 4.285 mg pitstop-2 (molecular weight of pitstop-2 is 428.5g/mol) in 1 mL of DMSO.

### 5.13 4% Paraformaldehyde

- Heat 440 ml of distilled water to 60 °C, ensuring it does not exceed 65 °C. If it fails, discard it and begin anew.
- Dissolve 20 g of paraformaldehyde in water and add approximately 50 µL of 10N NaOH.
- After dissolving PFA, add 50 mL 10X PBS, pH 7.4.
- Check the final pH; it should be 7.4, and the total volume should be 500 ml. Store at 4 °C.

### 5.14 DMEM: F12 (Lactose + pitstop-2) working solution

- Add 2.0 µL of 10 mM Pitstop-2 (DMSO) to a sterile tube and add 150 µL of 1 M lactose to the same tube.
- Add 848 µL pre-warmed serum-free DMEM: F12, gently mix by pipetting up/down.

Use immediately for treatment.

## 6 Synthesis Procedure

### 6.1 Glassware cleaning

- Place all necessary chemicals inside the fume hood. In a 250 mL beaker, carefully measure acetic acid and nitric acid in a 3:1 ratio to make an aqua regia solution.
- Transfer 80 mL of this solution into a 100 mL beaker, then dip the mouth of the condenser into it and seal the top with tissue paper.
- Pour the remaining 100 mL of aqua regia into a round-bottom flask (RBF) containing a magnetic stir bar. Seal the mouth with tissue paper. Leave both the condenser and RBF undisturbed for 24 hours.
- After 24 hours, discard the Aqua regia solution and ensure that it is completely neutralized before discarding.
- Wash the RBF and condenser thoroughly with soap solution and rinse sequentially with acetone and then Milli-Q water. Allow the glassware to air-dry for a few minutes.
- For drying and sterilization, cover the mouths of the RBF and condenser with aluminum foil. Using a sterile syringe needle, make small punctures in the foil to allow ventilation. Place the glassware in a hot air oven for 24 hours to achieve complete drying and sterilization.

### 6.2 Solution preparation

- Weigh 1-1 gm of citric acid and ascorbic acid, and dissolve in a 1:2 ratio of Ethanol and Milli-Q water, and mix them in a vortex mixer for 5 minutes. Use sonication for 10 minutes to ensure complete dissolution and homogeneity.

### 6.3 Reflux

- Gather all the required apparatus and glassware in the fume hood and turn on the fume hood.
- Place the setup on a magnetic stirrer with a temperature probe and place the oil bathtub on it.
- Pour the prepared reaction mixture into a round-bottom flask (RBF) containing a magnetic bead.
- Attach the condenser to the RBF and seal the joint securely with Teflon tape to prevent vapor leakage. Furthermore, insert a tissue paper plug at the open end (top) of the condenser.
- Connect the inlet and outlet tubes of the condenser to a water pump to ensure continuous cooling during reflux.
- Start the pump and add ice to the bucket and eventually change the ice every 30 minutes for 12 hours.
- Adjust the temperature to 130 °C, set the stirring speed to 300 rpm, and allow the system to run under reflux conditions for 12 hours.

### 6.4 Cooling & Collection

- After completion of 12 hours of reflux, gradually reduce the heating and allow the reaction mixture to cool down slowly to room temperature (≈27 °C).
- Transfer the cooled mixture into a 50 mL Falcon tube and wrap the Falcon tube with aluminum foil to protect the sample from light exposure. Store the Falcon tube under light-protected conditions until further use.

### 6.5 PH Maintenance

- Measure the pH of the GCQD solution, which is approximately 1.85-3. Add the NaOH solution dropwise into the GCQD solution while continuously stirring.
- Monitor the pH using a calibrated pH meter. Continue the addition until the solution reaches a neutral pH of 7.0.

### 6.6 Dialysis

- Measure the 15cm of dialysis bag and dip it in Milli-Q water in a 1000 mL beaker for 10-15 minutes.
- Now take out the dialysis bag from the water and remove excess water from each dialysis bag. Seal one end of the bag tightly with clips.
- Fill the sample solution into the dialysis bag from the open end and seal the open end with clips.
- Check for leakage by placing the ends on tissue paper and repeat the same process for all dialysis bags. Again, release the sample-loaded dialysis bags into the Milli-Q water-filled beaker.
- Place the beaker on a magnetic stirrer, start gentle stirring at room temperature for 24 hours.
- Replace the Milli-Q water after the first hour. Continue to replace the water at 6-hour intervals for the remainder of the dialysis period.
- After 24 hours, remove the dialysis bags from the beaker. Collect the dialyzed samples into Falcon tubes.

### 6.7 Lyophilization

- Divide the samples into appropriate volumes in Falcon tubes and remove the caps of the Falcon tubes. Furthermore, cover the openings with parafilm.
- Using a sterile syringe needle, make small punctures in the parafilm to allow vapor escape during freezing and drying.
- Place all Falcon tubes carefully in a −80 °C freezer and freeze for 24 hours.
- After freezing, collect the tubes and load them into the Lyophilizer for 24 hours.
- After lyophilization, collect the tubes containing the dry powdered form of GCQDs.

Store appropriately in airtight containers for further use, protected from light.

## 7 Characterization

### 7.1 UV-Vis spectroscopy

- Weigh 1 mg of the sample and dissolve it in 1 mL of Milli-Q water and mix thoroughly using a vortex mixer until the sample is completely dissolved.
- Clean and rinse the cuvettes with Milli-Q water to avoid contamination, and add 1 mL of the prepared sample solution into the cuvette.
- Place the cuvette inside the UV-Vis spectrophotometer chamber and record the absorbance spectrum in the wavelength range of 200-800 nm.
- Save the obtained spectra and process the image/data.
- Plot the absorbance data using Origin software to generate the final graph.

### 7.2 Fluorescence spectroscopy: -

- Weigh 1 mg of the sample and dissolve it in 1 mL of Milli-Q water inside a 2 mL microcentrifuge tube (MCT). Mix thoroughly using a vortex mixer until fully dissolved.
- Clean and rinse the cuvettes with Milli-Q water and pipette 1 mL of the prepared solution into a cuvette.
- Place the cuvette in the fluorescence spectrophotometer chamber.
- Set the excitation wavelength (commonly ∼450 nm for GCQDs) and record the emission spectra across the desired range (e.g., 480-650 nm).
- Save the fluorescence spectra and process the data using Origin software and plot the emission graph.

### 7.3 Fourier Transform Infrared Spectroscopy (FTIR)

- Take the Falcon tube containing the dried GCQDs sample and weigh 1 mg of the dry residue using a microbalance.
- Clean the sample holder/placement area of the FTIR instrument with isopropanol (IPA) and tissue paper to remove contaminants and place the dry sample onto the FTIR sample slot.
- Secure it by arranging the pressure tower and compression tips properly.
- Set the scanning range of the FTIR instrument to 4000-500 cm^-1^ and begin recording the spectrum.
- Save the acquired FTIR spectrum and process the data using Origin software to generate and analyze the graph.

### 7.4 X-ray diffraction (XRD)

- Clean the sample placement area of the XRD instrument to remove any dust or contaminants. Take 1 mg of the solid powdered sample and place the powdered sample evenly in the XRD sample holder.
- Run the XRD measurement and record the diffraction pattern, and save the acquired data. Process the diffraction pattern using appropriate software to generate the XRD graph.

### 7.5 Atomic Force Microscopy AFM

- Take 100 µL of the sample and dilute it with 900 µL Milli-Q water (1:10) in a PCR tube.
- Cut a small piece of mica sheet and fix it onto a glass slide using clear nail paint adhesive. Drop-cast 10 µL of the diluted sample onto the mica-coated glass slide.
- Place the prepared slide inside a desiccator for 10–15 minutes to allow drying, and mount the prepared glass slide on the AFM sample stage and align it properly.
- Attach and set the AFM cantilever, select and activate the tapping mode from the instrument software.
- Acquire AFM images, process and analyze the data using JPK software.

### 7.6 Transmission Electron Microscope (TEM)

- For sample preparation, we first have to dissolve the 1mg sample in 1 mL of Milli-Q water.
- Take 10µL from the dissolved sample and again dissolve in 1 mL Milli-Q water.
- Centrifuge and drop cast 5 µL on a TEM grid, place the TEM grid, process the Image, and save the data.

## 8 Quantum Yield Calculation

- To figure out the Quantum Yield of GCQDs, we have to take FITC as the reference.

For the Quantum Yield calculation, we need to prepare a solution of FITC at a concentration of 50 µg/mL in DMSO.

- The UV absorbance and fluorescence were quantified by adding 0.5 µL of FITC from the produced stock solution to the cuvette, achieving a final volume of 3 mL.
- Likewise, 1 µL of RB (Rhodamine B) was administered for the subsequent reading, maintaining a final volume of 3 mL. Additionally, the following measurements were obtained by maintaining the final volume at 3 mL while increasing the FITC concentration by 0.5 µL.
- To quantify the fluorescence and UV absorbance, 5 µL of GCQDs (0.2 mg/ml) were added to the cuvette, maintaining a final volume of 3 mL. Once more, the GCQD concentration was increased by 5 µL for the successive readings, while maintaining a final volume of 3 mL. Both FITC and GCQD absorbances were maintained below 0.1 a.u. (FITC from literature, Φ = 0.92).
- GCQDs were dissolved in water and have a refractive index of 1.33, and FITC is dissolved in DMSO and has a refractive index of 1.47. Their fluorescence spectra were recorded at the same excitation of 480 nm. Then, by comparing the integrated photoluminescence intensities (excited at 480 nm) and the absorbance values of GCQDs with the reference FITC, the quantum yield of the GCQDs was determined.
- The data is plotted (Figure 3), and the slopes of the sample (GCQDs) and the standard (FITC) were determined. The data showed good linearity, with a mean square deviation of 0.987 for the reference FITC and 0.994 for the sample GCQDs.
- The quantum yield was calculated using the given formula equation given below:

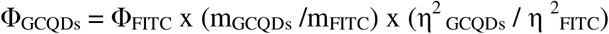
- Where Φ is the quantum yield, η is the refractive index of the solvent, m is the slope, FITC is the reference, and GCQDs is the sample. The quantum yield for GCQDs was found to be 0.055%.
- The quantum yield calculation of GCQDs was calculated by using rhodamine B (RB) as a reference. For quantum yield calculation, dissolve 1mg rhodamine in 1 mL Milli-Q (1mg/mL). further take 0.1mL from the 1mg/mL solution and dissolve in 9.9mL of milli-Q water. The final concentration is 10µL.
- The UV-absorbance and fluorescence spectroscopic data were taken by taking 4 µL of RB and keeping the final volume to 3mL. Similarly, the subsequent readings were taken by increasing the concentration of RB by 2 µL, keeping the final volume to 3mL.
- The UV-absorbance and fluorescence spectroscopic data of GCQDs were taken by taking 5µL of GCQDs in the cuvette, keeping the final volume to 3mL. the subsequent readings were taken by increasing the concentration of GCQDs by 5 µL, keeping the final volume to 3mL.
- The absorbance for both RB and GCQDs was kept less than 0.1 at 540 nm. RB (literature Φ = 0.31) and GCQDs were both dissolved in water (refractive index = 1.33).
- Comparison between the photoluminescence intensities (excited at 500 nm) and the absorbance values of GCQDs with the reference RB quantum yield should be determined, as shown in Figure 3, and the comparison of the slopes of the sample(GCQDs) and the standards (RB) was also determined. These characterized data show the linearity mean square deviation 0.99 (reference) and sample(0.95).
- For the Quantum yield calculation, we use the equation given below

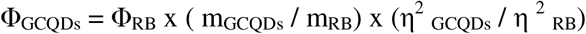
- Where Φ is the quantum yield, η is the refractive index of the solvent, m is the slope, RB is the reference, and GCQDs is the sample. According to the given equation, the quantum yield for GCQDs was calculated to be 3.3 %.

## 9 Cytotoxicity assay of GCQDs in RPE1 cells and tissue-derived primary cells

- To perform the cytotoxicity assay of GCQDs, firstly, we have to seed approximately 1×10^4^ per 100µL of RPE1 (Retinal pigment epithelial cell line) cells, and tissue-derived cells (liver and Kidney) cells.
- These cell lines were seeded in a 96-well plate, which already contains media. These media contain (DMEM containing 10% FBS, sodium bicarbonate, 1xAntibiotic (penstrep), HEPES, sodium pyruvate) and are allowed to accumulate for 24 hours in 5% CO_2_ at 37°C before the experiment.
- Check the cells the next day, and the cells were washed with PBS, and GCQDs treatment is given in increasing concentration of GCQDs (50,100,150,200,250,300,350,400µL/mL) for 24 hours in a serum-free medium. After exposure, the culture medium should be removed from the cells and discarded.
- Again, DMEM containing MTT salt (5 mg/mL) was added and further incubated for 3-4 hours for the determination of the mitochondrial dehydrogenase activity of viable cells.
- Again, the medium was removed, and 100 µl of DMSO was added to each well in 96 96-well plate to dissolve the purple formazan crystals and to measure the absorption spectrum by taking absorption at 570 nm using a Multiskan microplate reader.
- The experiment was done in triplicate, normalized to the corresponding well containing DMSO, whereas the non-treated GCQDs well was considered as a control to calculate the % cell viability check of each well.

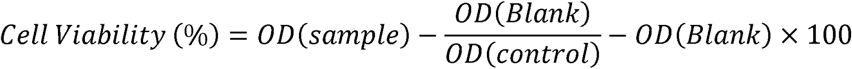

## 10 Preparation of mouse kidney, liver, heart, and brain primary cells

- 1-day-old neonatal mice were rinsed off with a 70% ethanol solution for complete surface sterilization and decapitated with sterilized scissors. The skin was cut, and the chest was opened along the sternum, and carefully expose the chest cavity.
- Now, immediately place extracted organs in sterile cold PBS (without Ca²L and Mg²L). wash with PBS and chop tissues into small pieces.
- Add 1x collagenase D solution and incubate for 20 minutes at 37°C.
- Pass digested tissue through a sterile cell stainer to obtain a single-cell suspension and centrifuge for 10 minutes at 2000 rpm with a maintained 4°C temperature.
- Collect the cell pellets and wash with PBS; furthermore, resuspend the DMEM: F12 (1:1) medium and seed in a culture flask, and allow the cells to grow until they reach confluence.

## 11 Cellular Uptake of GCQDs in kidney, liver, heart, and brain primary cells

- For the investigation of the GCQDs pathway into the cell, a clathrin-mediated (CME) and a clathrin-independent (CIE) pathway were performed in prepared primary cells isolated from the mice kidneys, liver, heart, and brain. Confluent cells were trypsinized and seeded in a 24-well plate on a coverslip at a density of 5 × 10^4^ cells.
- Positive marker treatment is given in which cells are incubated with transferrin (5µg/mL) as a clathrin-mediated endocytosis (CME) marker and Galectin-3 (5 µg/mL) as a clathrin-independent endocytosis (CIE) marker.
- Treatments were performed alone and in combination with GCQDs at increasing concentrations of 50, 100, and 200 µg/mL, with incubation in 5% CO_2_, at 37°C for 30 minutes, to investigate the CME and CIE pathways, respectively.
- To study the endocytic pathways involved in GCQDs internalization, GCQDs are incubated with the Pitstop-2 (20µM) as a CME inhibitor and lactose (150mM) as a CIE inhibitor.
- These cells were pre-incubated with inhibitors in serum-free media for 30 minutes at 37°C, 5% CO_2_.
- For the CME pathway, cells were first treated with pitstop-2 alone for 10 min. then, they were treated with pitstop-2 + GCQDs together to evaluate uptake reduction. The untreated cells served as blank controls.
- For further fixation and staining after treatment, cells are washed with 3× with PBS. Followed by Fixation with 4% paraformaldehyde (PFA) for 10 minutes at 37°C and again washed with 3× with PBS. Now add Coverslips mounted using Mowiol mounting solution containing Hoechst dye.
- Confocal microscopy was performed to visualize GCQDs uptake and pathway involvement.

## 12 Ex vivo Cellular Uptake of GCQDs in Mouse Kidney, Heart, and Liver Tissue Sections: -

- For tissue preparations, isolate the kidney, heart, and liver tissues from mice and wash the tissues with cold PBS. Snap-freeze immediately in liquid nitrogen and cut into 1mm slices using a sterile scalpel.
- For revival of tissue slices, transfer the frozen tissue slices into DMEM: F12 (1:1) medium and incubate for 20 minutes at 37°C, 5% CO_2_.
- For pretreatment wash, tissue slices are once washed with PBS for 5 minutes and cut using a sterile scalpel.
- For GCQDs treatment, these slices were incubated with GCQDs at different increasing concentrations of 50, 100, and 200µg/mL with serum-free DMEM: F12 medium for the duration of 60 minutes at 37°C,5%CO_2_.
- For washing and fixation, we wash the slices 3× with PBS to remove unbound GCQDs and fixed with 4% paraformaldehyde (PFA) for 15 min at 37 °C, with further washing with PBS.
- Now mount slices using Mowiol mounting solution containing Hoechst dye, and ensure complete mounting.
- Perform confocal microscopy to study ex vivo uptake of GCQDs.

## 13 In vivo uptake of GCQDs in Zebrafish Model: -

- In vivo uptake assays were performed according to the Organization for Economic Cooperation and Development (OECD) guidelines.
- For the larvae preparation, at 72 hours post-fertilization (hpf), remove dead larvae and transfer the live larvae into six-well plates (15 larvae per well).
- Provide the GCQDs treatment to the larvae at two concentrations, the first is 100µg/mL and the second is 200 µg/mL. One well in each group was kept as a control (without nanoparticles) and incubated for 6 hours.
- For post-treatment processing, replace the medium with fresh E3 medium and wash larvae thoroughly to remove excess GCQDs.
- For further fixation, 4% Paraformaldehyde (PFA) for 2 minutes is used to fix the larvae.
- Mount larvae with Mowiol mounting solution, allow to dry, and perform confocal microscopy for uptake analysis.

## 14 Confocal Imaging and Processing

- The confocal imaging of fixed cells (63x oil immersion) and fixed tissues/embryos (10x) was performed using Leica TCS SP8 confocal laser scanning microscope (CLSM, Leica Microsystem, Germany).
- Different fluorophores were excited with different lasers, i.e., for Hoechst (405 nm), GCQDs (488 nm), Tf (633 nm), and Gal (633 nm). The pinhole was kept 1 airy unit during imaging.
- Image quantification analysis was performed using Fiji ImageJ software. For the quantification analysis, whole cell intensity was measured at maximum intensity projection, background was subtracted, and the measured fluorescence intensity was normalized against unlabelled cells.
- A total of 40-50 cells were quantified from collected z-stacks for each experimental condition.

## 15 Results and Discussions

We prepared GCQDs with biomolecules, citric acid (CA), and Ascorbic acid (AA) in solution with water and ethanol (2:1) ratio. (Figure 1) shows that the dissolved precursor was left to reflux at 130°C in the solvent over a period of 12 hours. NaOH solution is added to the pH till it attains 7 pH. GCQDs were purified by 24 hours of dialysis with Milli-Q water. Additionally, we dried the GCQDs using lyophilization to a powdery sample. The GCQDs obtained by the synthesis process were thoroughly characterized with several biophysical and spectroscopic techniques, which are presented in Figure 1. The absorption spectrum of the GCQDs in water (pH 7) is illustrated in UV-visible Spectroscopy with two characteristic peaks at 265 nm and 365nm. The n-π* transition of the C = O bond Figure 2a is found at 265 and 365nm. (Figure 2b) shows that the photoluminescence (PL) of GCQDs was highly emitted under 450nm excitation with a maximum intensity of 525nm. The FTIR spectra of the GCQDs have verified the existence of a number of surface functional groups. A wide band of 3232 cm^-1^ stretch vibration was associated with the O-H stretching vibrations, and a peak at 2989 cm^-1^ indicated sp^3^ C-H stretching. The signal at 1722 cm^-1^ was ascribed to carbonyl (C=O) vibrations, and the signal at 1566 cm^-1^ indicated C=C vibrations. This was further supported by other peaks at 1390 cm^-1^(sp^3^ C-H bending), 1262 cm^-1^(C-O stretching), and 1045 cm^-1^(C-O-C stretching) to show the functional diversity on the GCQD surface. In comparison to the predecessors of citric acid and ascorbic acid, the appearance of the peak shifts and new signals proved the successful formation of a bond and surface modification. These chemical characteristics, especially the hydroxyl and carboxyl groups, enhance the water solubility and dispersibility of the synthesized GCQDs (Figure 2b). (Figure 2d,e) showed that the GCQDs had a morphology size of 23 nm and a spherical shape as seen under transmission electron microscopy (TEM). Broad peaks at 11.32°, 21.89°, and 33.2° (2θ) in powder XRD indicated a largely amorphous structure (Figure 2f). The geometric properties of the GCQDs appear to be of a spherical nature and are depicted by atomic force microscopy (AFM). The GCQDs have a size and height of 2-3 nm, with the width of the majority of the particles being under 10 nm, as depicted in (Figure 2 g-i).

**Figure 1:**
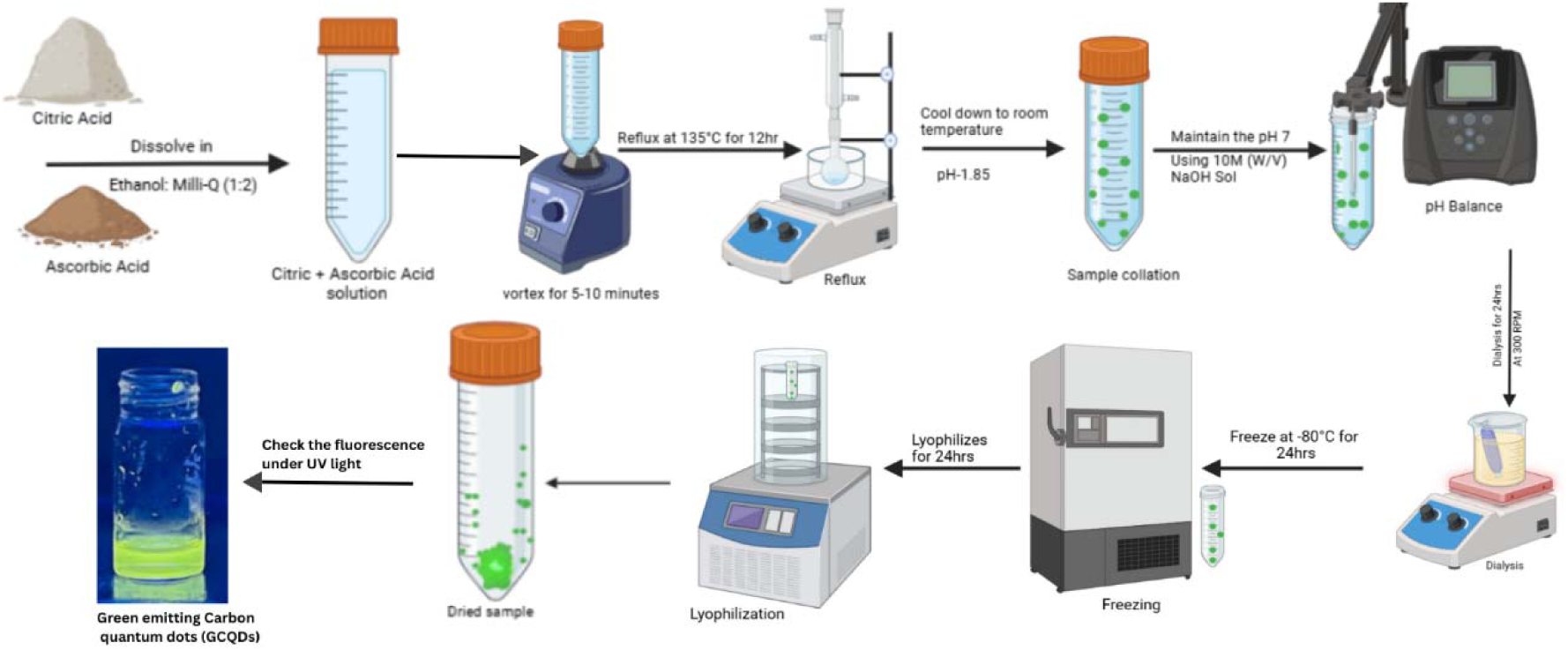
Illustrates the stepwise synthesis of green-emitting carbon quantum dots (GCQDs) using citric acid and ascorbic acid as precursors. Initially, the acids were dissolved in an ethanol: Milli-Q mixture (1:2) and vortexed for 5-10 minutes to obtain a homogeneous solution. The mixture was then subjected to thermal carbonization by refluxing at 135 °C for 12 hours, followed by cooling to room temperature, yielding a crude solution with a pH of 1.85. The pH was subsequently adjusted to 7 using 10 M NaOH, and the neutralized solution was purified by dialysis for 24 hours under constant stirring at 300 rpm. The dialyzed sample was frozen at −80 °C for 24 hours and further lyophilized for 24 hours to obtain a dry powdered form of GCQDs. The final product was collected in airtight containers and verified for green fluorescence under UV light.

**Figure 2:**
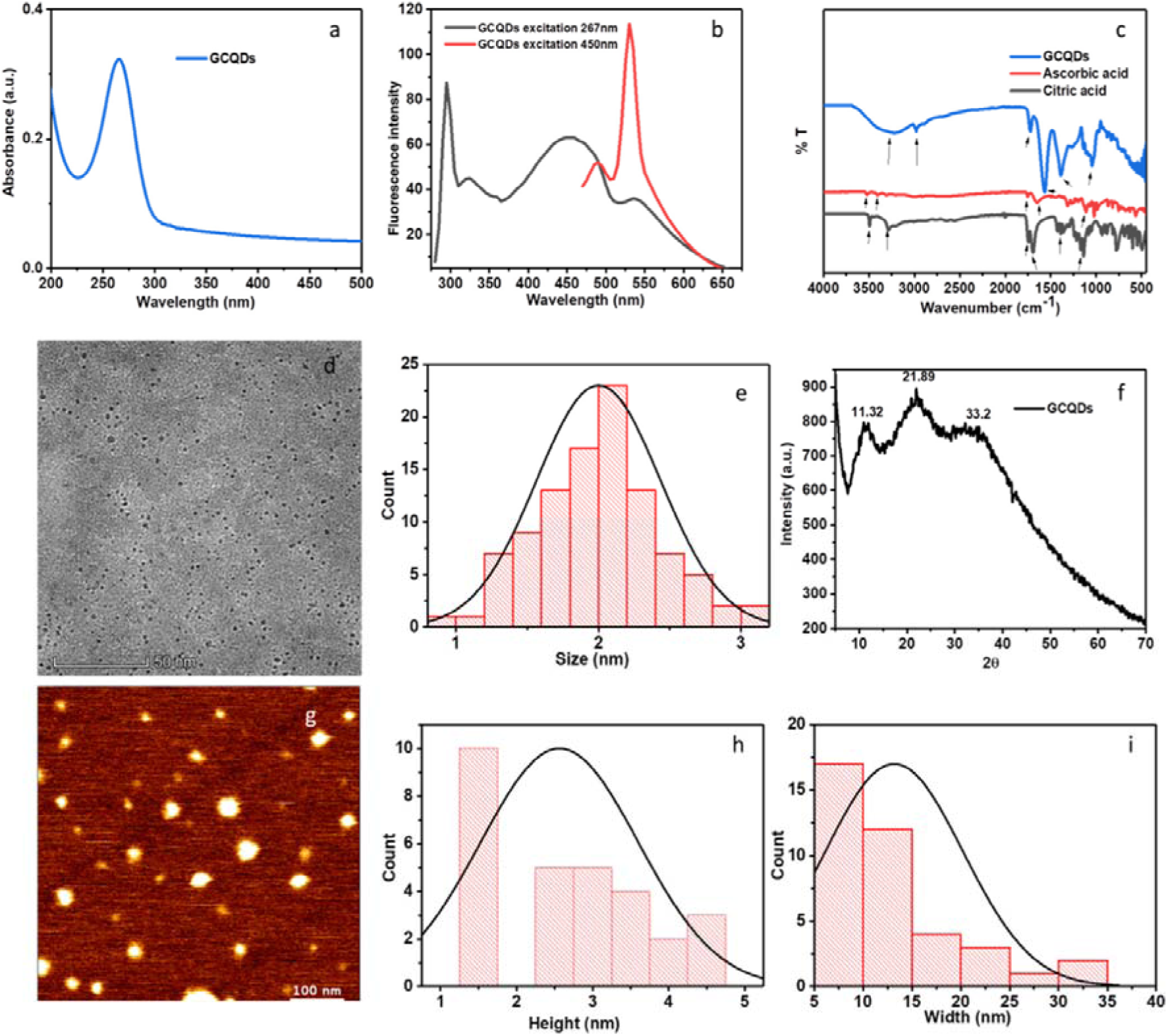
UV-vis spectroscopy of GCQDs dispersed in deionized water (pH 7) revealed a distinct absorption peak at 265 nm(Figure a). The photoluminescence spectra showed multiple emission bands at 300, 325, 450, and 525 nm upon excitation at 250 nm, with the strongest emission detected at 525 nm when the excitation wavelength was set to 450 nm (Figure b). FTIR analysis confirmed the presence of functional groups on the GCQD surface. A broad band at 3232 cmLJ¹ corresponded to O–H stretching vibrations, while the peak at 2981 cmLJ¹ represented C-H stretching. The signal at 1722 cmLJ¹ indicated carbonyl (C=O) groups, the band at 1390 cmLJ¹ corresponded to O-H bending, and peaks at 1262 cmLJ¹ and 1045 cmL¹ were assigned to C-O and C-O-C stretching, respectively. When compared with spectra of citric acid and ascorbic acid precursors, GCQDs exhibited shifted or missing peaks, for instance, the bands at 1744 cm^-1^ and 1992 cmL¹ (citric acid) and 1111 cm^-1^, 743 cmL¹, and 3525 cmL¹ (ascorbic acid) were absent in GCQDs, highlighting successful transformation of the precursors (Figure c). Transmission Electron Microscopy (TEM) imaging demonstrated that the particles were spherical and crystalline in nature (Figure d). A size distribution analysis based on 100 TEM-counted particles indicated dimensions in the range of 1-3 nm (Figure e). Powder XRD of lyophilized GCQDs (pH 7) showed broad peaks at 2θ = 11.32°, 21.89°, and 33.2°, confirming the largely amorphous structure of the material (Figure f). AFM analysis further validated the nanoscale morphology. Height profiles showed a thickness of 2-5 nm (30 particles measured), while width distribution analysis revealed that most particles had lateral dimensions of below 10 nm (Figure g-i)^35^.

### 5.1 Quantum Yield calculations data

The optical behavior of GCQDs was compared against established references systematically to ensure that it met the required optical fluorescence and could be used to estimate the quantum yield. As demonstrated in (Figure 3a), the GCQDs displayed a strong fluorescence signal in the aqueous medium when the GCQDs were excited at 500 nm, which confirms that it has a high intrinsic photoluminescence and stability in a biologically active solvent. The pattern of this intensity of these GCQDs indicates the efficient electronic transition and their capability of emitting in the visible range, which makes them applicable in bioimaging. To benchmark, (Figure 3b) shows the fact that Rhodamine B, a standard with clearly defined photophysical characteristics, will have an identical emission profile under the same conditions of excitation (500 nm) and solvent (water) as shown in (Figure 3b). The relative intensity of the emission patterns of GCQDs and Rhodamine B can be comparatively determined, which enables a relative evaluation of fluorescence efficiency and is calculated in quantum yields. To further assess the excitation-dependent behavior of GCQDs, Figure 3c depicts the fluorescence emission of GCQDs under the influence of excitation of the sample at 480 nm in aqueous solution. There was a constant emission, which indicates that GCQDs do not lose their strong fluorescence even at a slightly shorter excitation wavelength, which corresponds to their wide excitation wavelength and structural stability. Lastly, Figure 3d shows the emission response of fluorescein isothiocyanate (FITC) in DMSO, which is a popular reference, and which is excited at 480 nm. The use of FITC is a reliable comparison at 480 nm excitation and, at the same time, confirms that GCQD measurements are reproducible between solvents and reference standards. Altogether, all these data prove that GCQDs have stable and excitation-dependent fluorescence characteristics comparable with the reference groups like Rhodamine B and FITC. Their dark colors in aqueous solutions, specifically, make them especially attractive as far as bio-related applications are concerned, in terms of water solubility and photostability being of utmost importance.

**Figure 3:**
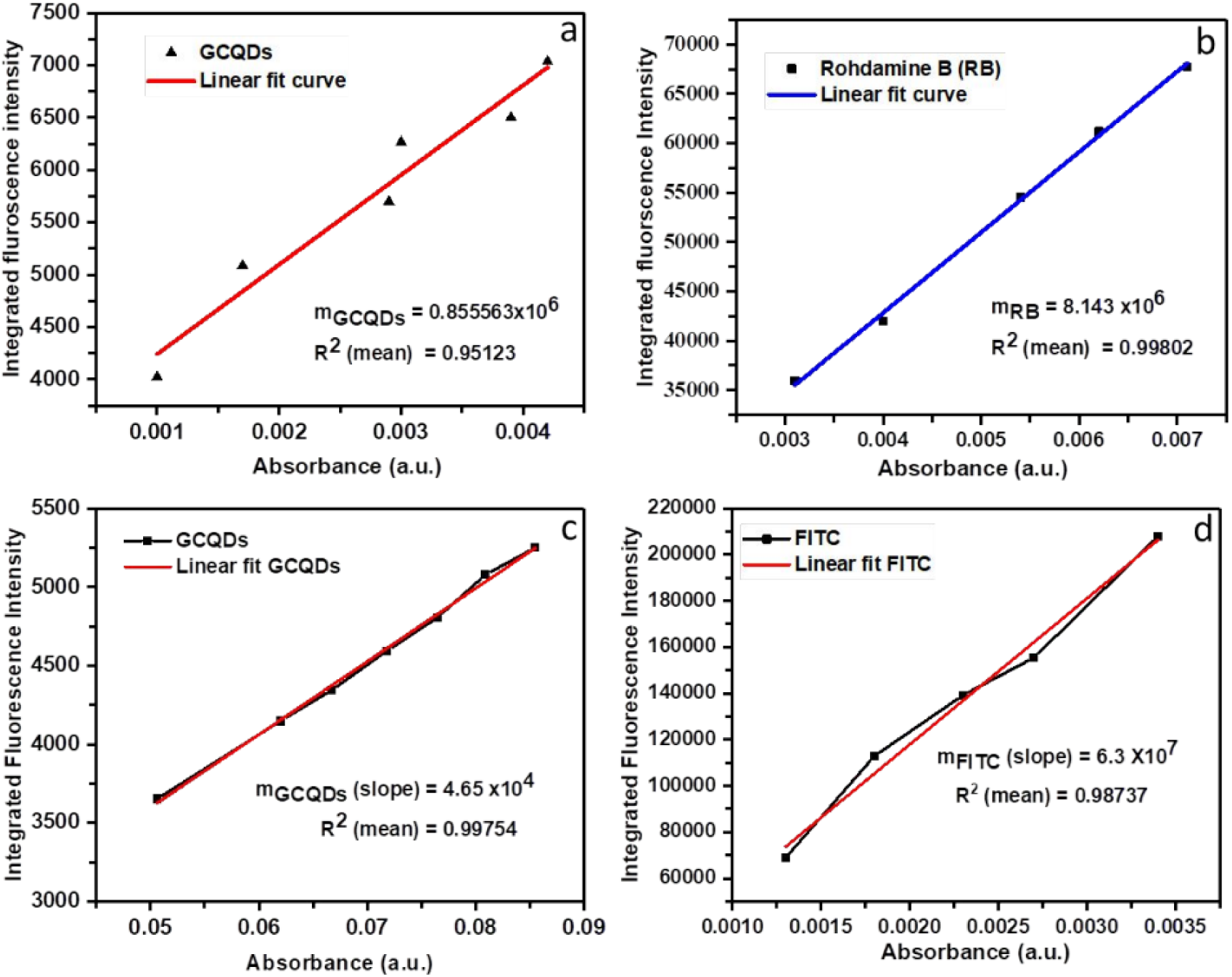
The first Plot (a) shows the total fluorescence intensity of GCQDs recorded upon excitation at 500 nm in aqueous medium. The second Plot (b) shows us the comparative plot of the total fluorescence response of Rhodamine B, used as a standard reference, also excited at 500 nm in water solvent. The third plot(c) displays the cumulative fluorescence intensity of GCQDs under excitation at 480 nm in water, and the fourth and last one shows (d) the corresponding plot of the fluorescence emission of FITC, taken as another reference dye, excited at 480 nm in DMSO solvent^35^.

### 15.2 Cellular Internalization and Nontoxicity of GCQDs: -

In order to investigate how GCQDs are internalized, mouse kidney, liver, heart, and brain primary cells were examined with clathrin-mediated endocytosis (CME) and clathrin-independent endocytosis (CIE) endocytosis studies. Pathway-specific inhibitors of CME (Pitstop-2, 20mM) and CIE (lactose, 150mM) were used to verify pathway participation, while transferrin (5 µg/mL) and galactin-3 (5 µg/mL) were used as positive markers of CME and CIE, respectively. Cells were incubated with GCQDs at different concentrations (50, 100, and 200 µg/mL), in the absence of markers or after adding markers and inhibitors. This was followed by confocal microscopy, used to visualize uptake, and then an assessment of fluorescence intensity was done. Transferrin uptake into kidney and liver cells was evident in both (Figures 4a and 4e), though its internalization was highly suppressed by pretreatment using the clathrin-specific inhibitor Pitstop-2 (20 µM). On the same note, the competitive inhibitor lactose (150 mM) did not result in a significant decrease in transferrin uptake, thus confirming the specificity of Pitstop-2 to prevent CME. In line with that, (Figures 4b and 4f) show the uptake profile of GCQDs in the same cellular models with inhibitors. The extent of internalization with GCQD was greatly reduced with the treatment with Pitstop-2, and there were no meaningful changes with the treatment with lactose. This tendency is a strong indication that in these cells, CME and not CIE is the main mechanism of uptake of GCQDs. These observations were also confirmed by quantitative analysis. The normalized fluorescence intensity of transferrin in the kidney and liver cell (Figure 4c and 4g) demonstrates that the Pitstop-2 treatment caused a statistically significant decrease (p < 0.0001), and the lactose treatment did not significantly affect it (ns). Another trend was noted in GCQD uptake quantification (Figure 4d and 4h), whereby Pitstop-2 strongly reduced fluorescence signals of internalised GCQDs, which supports the hypothesis of CME-mediated internalisation. Taken together, these findings prove that clathrin-mediated endocytosis is the most common entry route of GCQD into primary kidney and liver cells. The phenomena of co-localization with transferrin and the potent inhibitory effect on Pitstop-2 give strong mechanistic support to the use of CME in GCQD uptake. These findings play a vital role in the intracellular trafficking of GCQDs and the role of CME in determining their biodistribution and cellular fate.

**Figure 4:**
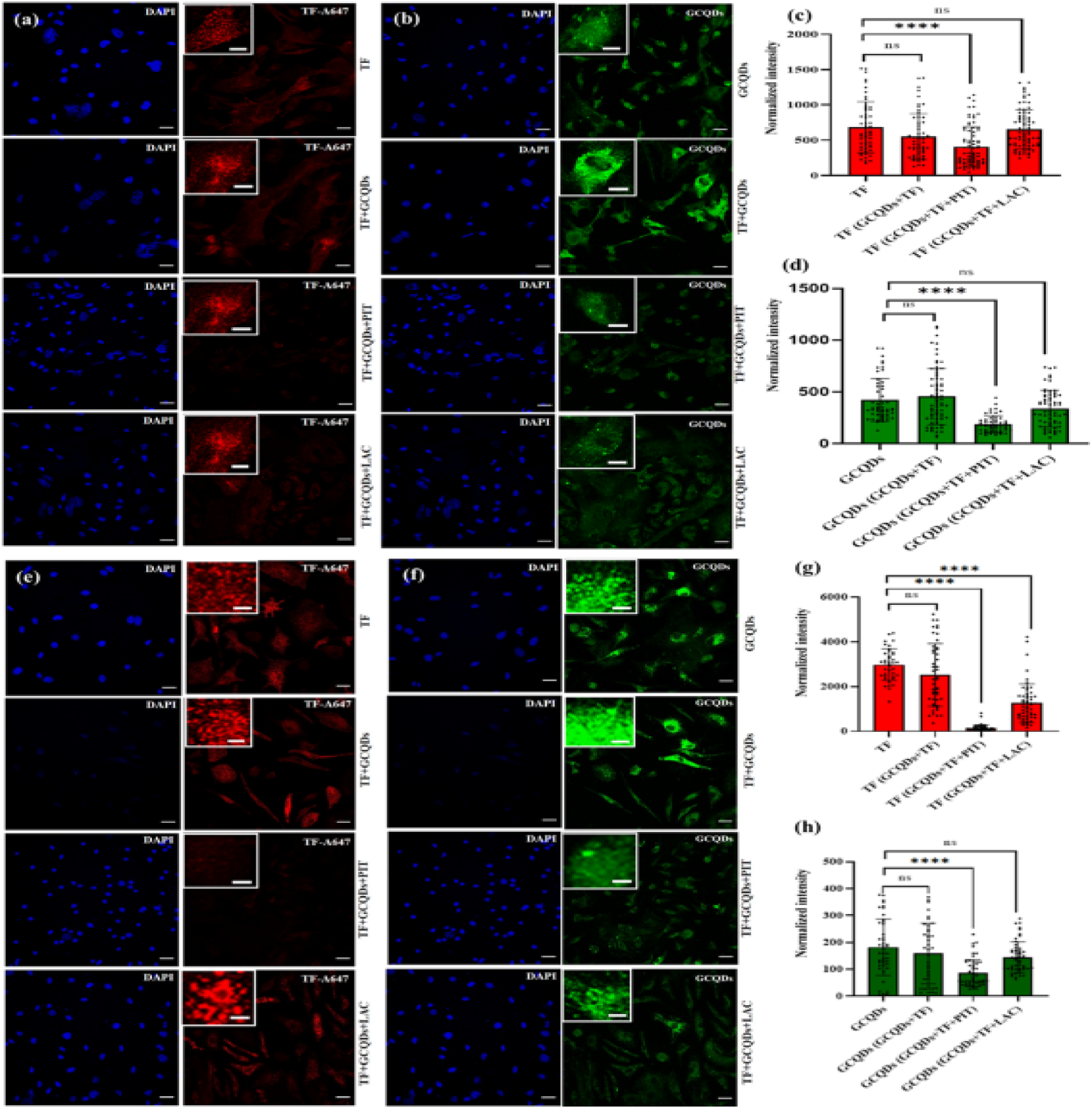
Clathrin-mediated endocytosis of transferrin (Tf) and GCQDs in mouse kidney and liver primary cells. (a, e) Internalization of A647-labeled transferrin (Tf) in kidney and liver cells following treatment with pitstop-2 (20 µM) and lactose (150 mM) in the presence of GCQDs. (b, f) Uptake of GCQDs in kidney and liver cells exposed to pitstop-2 (20 µM) and lactose (150 mM) in the presence of transferrin. (c, g) Quantification and normalized fluorescence intensity of transferrin (Tf). (d, h) Quantification and normalized fluorescence intensity of GCQDs. Scale bars: 20 µm (main images) and 0.75 µm (inset images). **** indicates a highly significant difference (p < 0.0001), while *ns* denotes no significant difference (One-way ANOVA). Data represent analysis of 50 cells from *n* = 2 independent experiments^35^.

To test the role of clathrin-independent endocytosis (CIE) in GCQD uptake, a CIE marker, Galectin-3 (Gal) was expressed in primary kidney and liver cells. A647-tagged Galectin-3 was incubated with cells in the presence of GCQDs, and some pathway-blocking agents were used: Pitstop-2 (20 µM) to inhibit CME and lactose (150 mM) to inhibit CIE. The control conditions, as demonstrated in (Figure 5a, e), indicated that Galectin-3 was strongly internalized, but the effect was significantly diminished after lactose treatment, which confirms its value as a dependable CIE marker. However, Galectin-3 uptake was not affected by Pitstop-2 treatment, suggesting that no clathrin-mediated processes contribute to Galectin-3 traffic. Parallel to this, GCQDs had uptake patterns comparable to Galectin-3 (Figure 5b, f). The treatment of lactose had a great impact in reducing internalization of GCQDs (****p < 0.0001), which implies that GCQDs also enter the cell by using CIE pathways. Interestingly, even a small but statistically significant decrease in CIE to their uptake was further impressed by a modest but statistically significant decrease in the number of certain conditions as well (p = 0.0287). These trends were always consistent in quantitative analyses (Figure 5c-h) but not in Pitstop-2 treatment (ns), supporting the pathway-specific inhibition. These results prove that, besides the clathrin-mediated endocytosis, GCQDs use clathrin-independent endocytosis pathways. The similarity in the behavior of GCQDs with Galectin-3 indicates the versatility of the uptake pathways of GCQDs, indicating that GCQDs have a dual internalization pathway. This versatility can increase the capacity to traverse cellular barriers, and this could boost their prospects in drug delivery, targeted bioimaging, and as a form of therapy.

**Figure 5:**
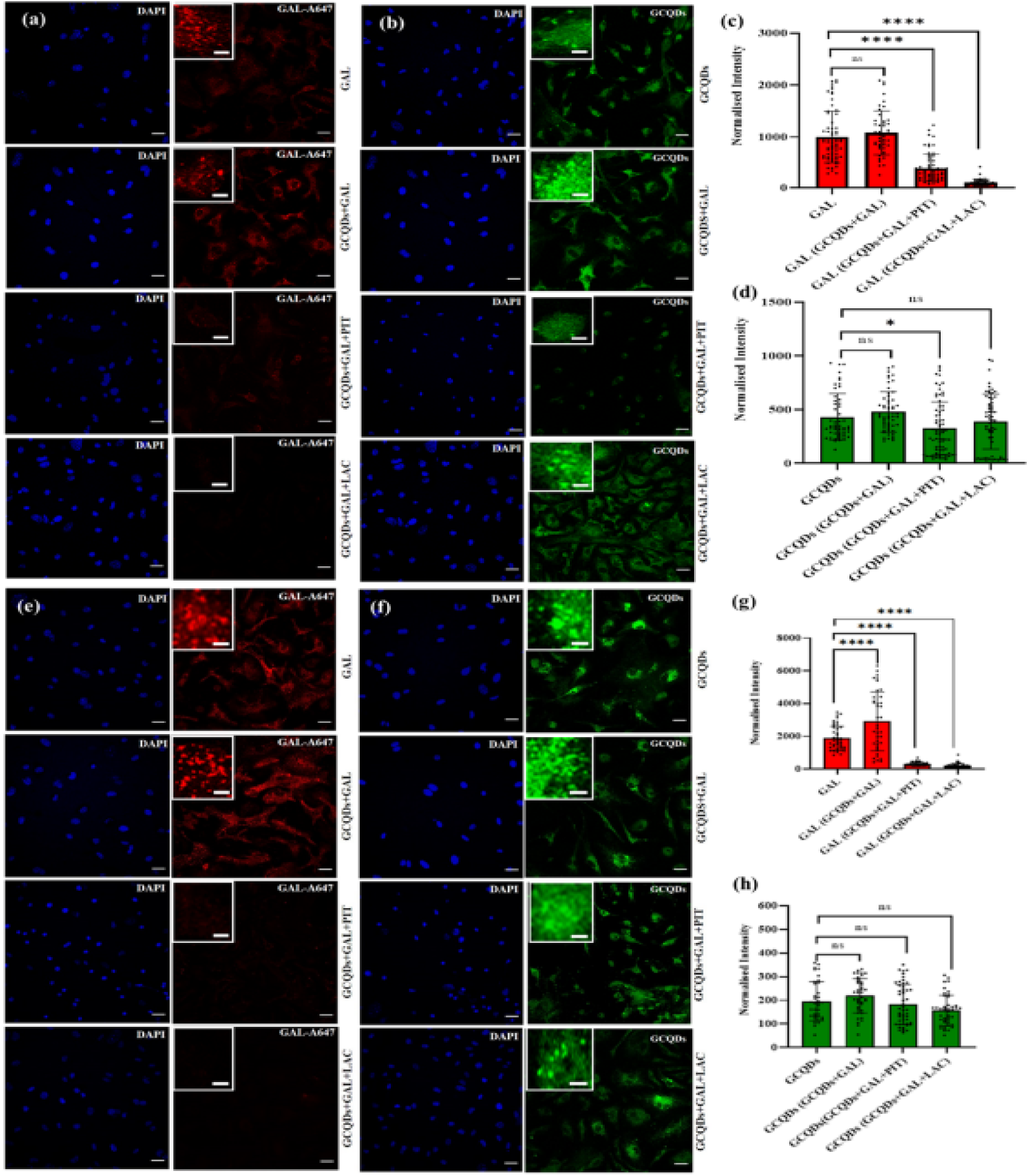
Endocytic uptake of galectin-3 (Gal) and GCQDs in mouse kidney and liver primary cells. (a, e) Internalization of A647-labeled galectin-3 in kidney and liver cells treated with pitstop-2 (20 µM) and lactose (150 mM) in the presence of GCQDs. (b, f) Uptake of GCQDs in kidney and liver cells exposed to pitstop-2 (20 µM) and lactose (150 mM) along with galectin-3. (c, g) Quantification and normalized fluorescence intensity of galectin-3. (d, h) Quantification and normalized fluorescence intensity of GCQDs. Scale bars: 20 µm (main images) and 0.75 µm (inset images). **** denotes a highly significant difference (p < 0.0001), *p = 0.0287 indicates a significant difference, and *ns* shows no significant difference (One-way ANOVA). Data represent analysis of 50 cells from *n* = 2 independent experiments^35^.

### 15.3 Ex Vivo and In Vivo Uptake Study of GCQDs in Mice-Derived Tissue Sections and Zebrafish Embryos

In order to compare the cellular uptake under a more physiologically relevant system, an ex vivo study is conducted using ex vivo kidney tissue sections, which were incubated with high concentrations of GCQDs and imaged under fluorescence microscopy. (Figure 6a) showed that GCQDs exhibited dose-dependent uptake in renal tissue versus untreated controls, and the higher the GCQD concentration in cells, the higher the intracellular fluorescence. The additional quantitative evaluation of the intensity of the fluorescence of the GCQDs (Figure 6b) disclosed that a considerable improvement in GCQD internalization with species increase. It was statistically significant (*p = 0.0326) at moderate concentrations, and it led to higher accumulation with higher concentrations (***p = 0.0004). These findings confirm a consistent finding that GCQDs not only interact efficiently with the isolated primary cells but also intravasate normal kidney tissue architecture. This in vivo study demonstrates the ability of GCQDs to undergo uptake into a tissue-relevant environment, which validates their potential across multiple fields in a translational approach to renal bioimaging and as therapeutic vectors. Additionally, its accumulation is concentration-dependent, implying that the uptake can be dosage-controlled, which will be a crucial parameter in the future optimization of biomedical use. To further understand the role of GCQDs in biodistribution in vivo beyond the in vitro and ex vivo models, we examined their in vivo uptake in zebrafish larvae, which is quite an accepted vertebrate model in nanomaterial toxicity and imaging investigations. After 6 or 12-hour exposure to 100 µg/mL and 200 µg/mL GCQDs, larvae had obvious fluorescence accumulation, and the accumulation was compared to the control sample (Figure 7a). Under the experimental conditions, brightfield images showed that the normal morphology of larvae was not caused by GCQD treatment (Figure 7a-d). Confocal imaging had demonstrated a powerful, concentration- and time-dependent uptake of GCQDs with a measurable fluorescence as early as 6 hours post-treatment (Figure 7e-g) but improved upon after 12 hours (Figure 7h). Localized GCQD accumulation was also seen in zoomed images (Figure 7i-l) that may indicate local uptake by the tissue. The statistically significant increase in uptake at both concentrations was confirmed in quantitative fluorescence analysis (Figure 7m), in which the effect was most significant at higher dose and longer exposure (****p < 0.0001; *p < 0.05). These findings confirm that GCQDs can penetrate into tissues of zebrafish larvae effectively and accumulate within the tissues in a dose- and time-dependent fashion, which justifies its possible use as bioimaging probes. Notably, adherence to OECD guidelines makes such findings have translational significance to nanotoxicological safety evaluations. The non-observation of clear toxicity throughout the exposure period also indicates the biocompatibility of GCQDs, but further research needs to be conducted on the longer-term clearance and organ-specific biodistribution^35^.

**Figure 6:**
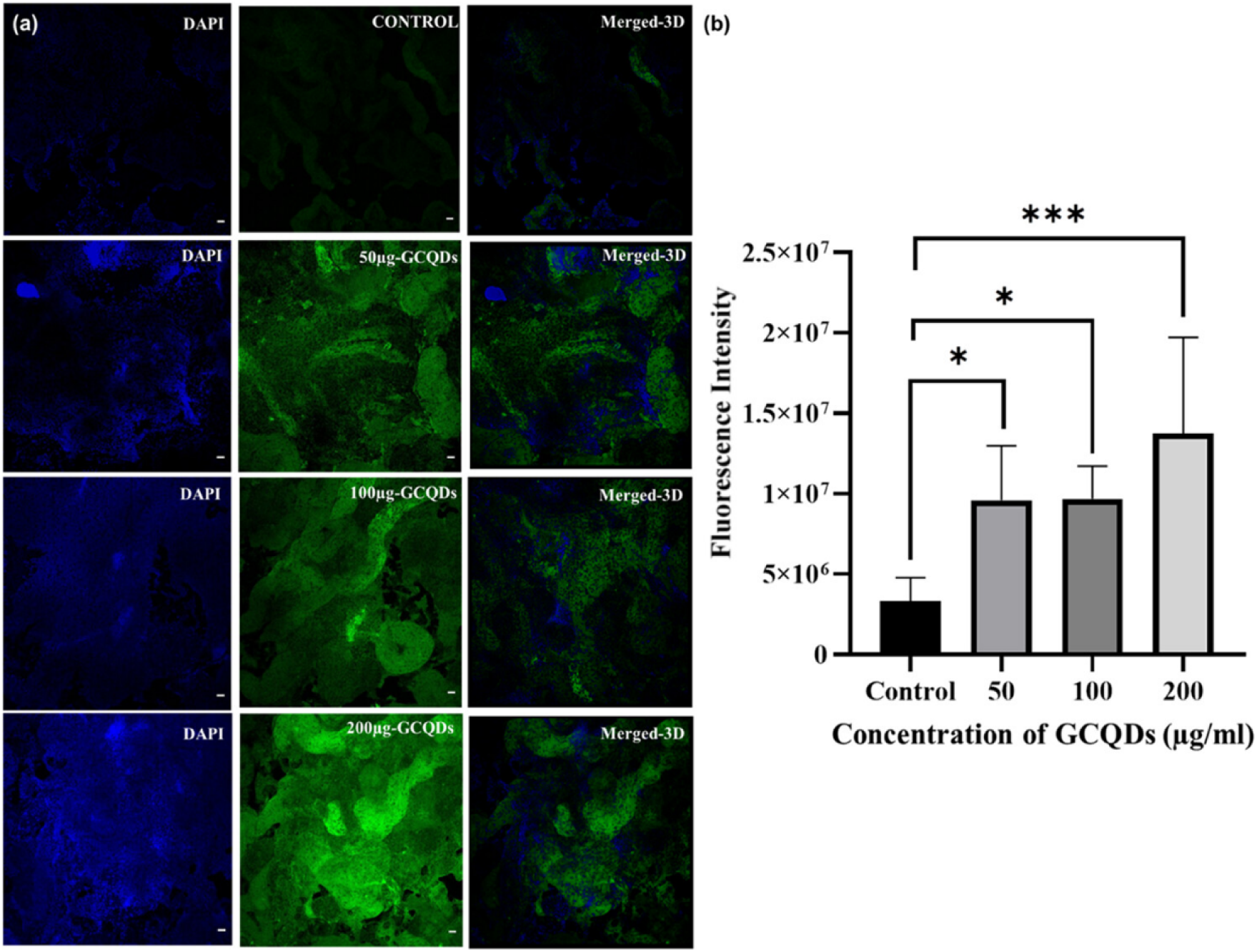
Uptake of GCQDs in kidney tissue. (a) Uptake of different concentrations of GCQDs vs nontreated GCQDs in kidney tissue sections. (b) Fluorescence intensity and quantitative measurement of the uptake of different concentrations of GCQDs in renal tissue slices. The scale bar is set at 20 μm. ***Statistically significant p-value (p = 0.0004). *Scale significantly significant p-value (p = 0.0326). n = 5 tissue slices per concentration were evaluated^35^.

**Figure 7:**
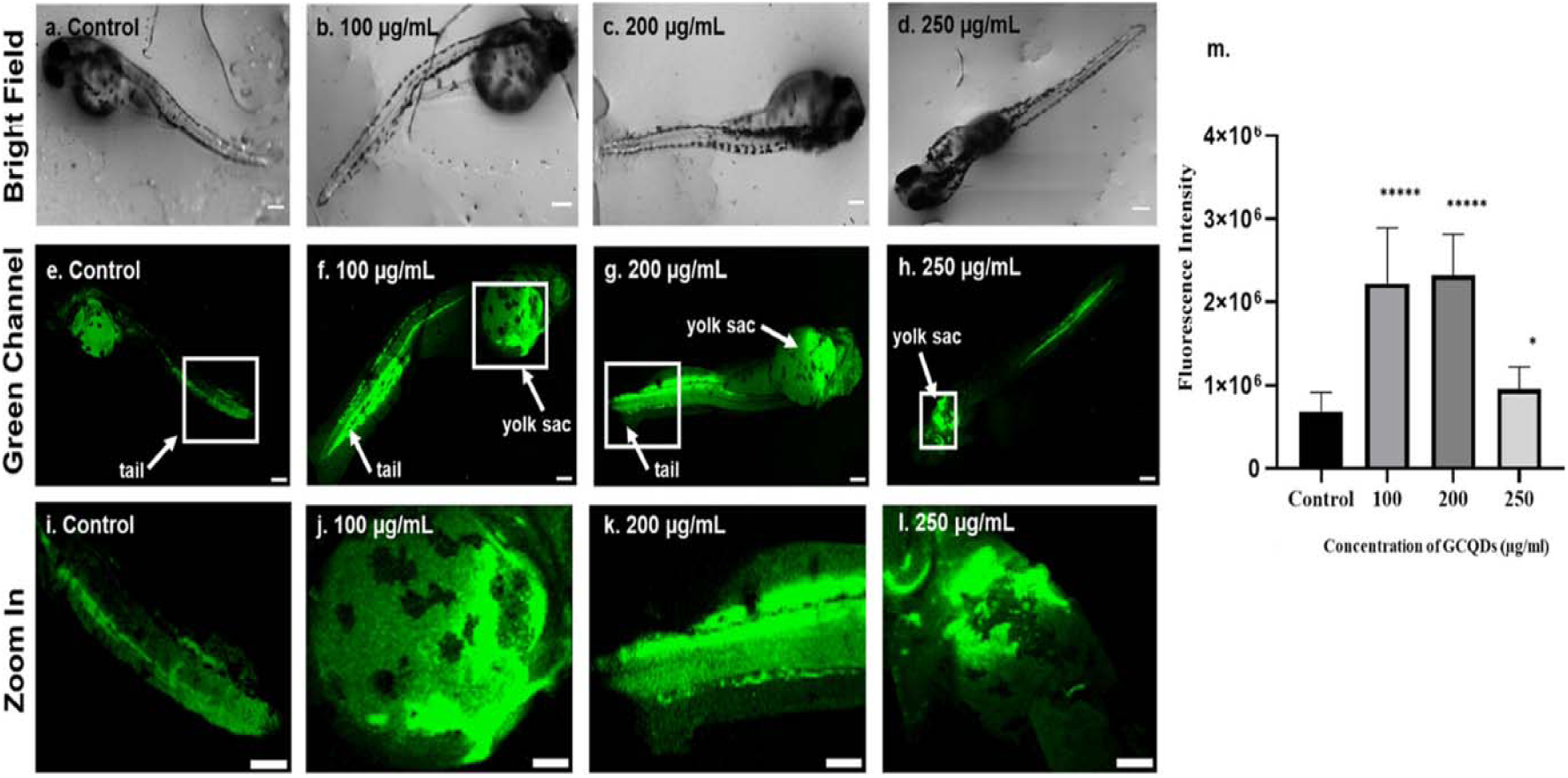
Uptake of GCQDs in zebrafish larvae. (7a) Uptake of different concentrations of GCQDs in 72 hpf zebrafish larvae following 6 and 12 h treatments. (a-d) Brightfield pictures of 72 hpf zebrafish larvae: (7e-g) 6 h treatment and (h) 12 h treatment. (7i-l) Zoom-in part of accumulated GCQDs in 72 hpf zebrafish larvae after 6 and 12 h treatments. (7m) Fluorescence intensity and quantitative study of the absorption of different concentrations of GCQDs in 72 hpf zebrafish larvae. The scale bar is set at 50 μm. ****Statistically significant p-value. (p < 0.0001). *Statistically significant p-value (p < 0.05). 08 larvae per condition were quantified^35^.

## 16 Conclusion

In this study, we successfully synthesized green carbon quantum dots (GCQDs) using precursor materials citric acid and ascorbic acid under reflux conditions, followed by a purification and lyophilization process to get a dry product yield. Further, we performed Comprehensive characterization by UV-Vis spectroscopy, Fluorescence spectroscopy, Fourier-transform infrared spectroscopy (FTIR), transmission electron microscopy (TEM), atomic force microscopy (AFM), and X-ray diffraction (XRD) confirmed their spherical morphology structure, nanoscale size distribution, amorphous structure, and abundant surface functional groups that imparted high water solubility and dispersibility. Quantum yield (QY) study and calculations, we use Rhodamine B and FITC as standards, demonstrating strong intrinsic fluorescence and photostability with excitation-dependent emission, making them suitable for optical and imaging applications. Furthermore, mechanistic uptake studies revealed that GCQDs predominantly enter primary kidney and liver cells through clathrin-mediated endocytosis, with additional contributions from clathrin-independent pathways, thereby these contributions highlighting their versatile internalization mechanisms. Ex vivo renal tissue analysis and in vivo zebrafish models further validated their efficient cellular penetration, dose-dependent accumulation, and non-toxic behavior. Collectively, all these studies demonstrate that GCQDs have excellent fluorescence properties, biocompatibility, and cellular uptake efficiency, establishing their potential as multifunctional probes for bioimaging, drug delivery, and future translational biomedical applications.

## Data availability

Source data for the figures in this protocol are available in the listed supporting primary research articles ^35^. Further parameters or details of the experiments are available from the corresponding author upon reasonable request.

## Supporting information

https://doi.org/10.1021/acsabm.3c00072

